# Short and Long-read Sequencing Survey of the Dynamic Transcriptomes of African Swine Fever Virus and its Host

**DOI:** 10.1101/2020.02.27.967695

**Authors:** Ferenc Olasz, Dóra Tombácz, Gábor Torma, Zsolt Csabai, Norbert Moldován, Ákos Dörmő, István Prazsák, István Mészáros, Tibor Magyar, Vivien Tamás, Zoltán Zádori, Zsolt Boldogkői

**Author notes:** corresponding authors: Zsolt Boldogkői, Dóra Tombácz.

## Abstract

African swine fever virus (ASFV) is an important animal pathogen causing substantial economic losses in the swine industry globally. At present, little is known about the molecular biology of ASFV, including its transcriptome organization. In this study, we applied cutting-edge sequencing approaches, namely the Illumina short-read sequencing (SRS) and the Oxford Nanopore Technologies long-read sequencing (LRS) techniques, together with several library preparation chemistries to analyze the ASFV dynamic transcriptome. SRS can generate a large amount of high-precision sequencing reads, but it is inefficient for identifying long RNA molecules, transcript isoforms and overlapping transcripts. LRS can overcome these limitations, but this approach also has shortcomings, such as its high error rate and the low coverage. Amplification-based LRS techniques produce relatively high read counts but also high levels of spurious transcripts, whereas the non-amplified cDNA and direct RNA sequencing techniques are more precise but achieve lower throughput. The drawbacks of the various technologies can be circumvented by the combined use of these approaches.

## Background & Summary

Methodological breakthroughs in sequencing technologies have revolutionized transcriptome profiling in recent years. Currently, the next-generation short-read sequencing (SRS) and third-generation long-read sequencing (LRS) platforms are widely used in genome and transcriptome research. SRS can generate large numbers of sequencing reads with unprecedented speed; however, it cannot sufficiently cover high-complexity transcriptomes. LRS produces lower data coverage with a higher error rate, but it can overcome many of the drawbacks of SRS, including the inefficiency in distinguishing between transcription isoforms and identifying embedded and long transcripts. The combined use of these platforms and library preparation chemistries can generate high-quality and throughput data on full-length transcripts. LRS has been applied for the assembly of transcriptomic maps in several organisms^1–8^, including viruses^9–17^.

African swine fever is a highly lethal animal disease affecting pigs and wild boars. The causative agent of this disease is the large, double-stranded DNA virus, the African swine fever virus (ASFV), the only member of the *Asfarviridae* family^18^. Because no effective vaccination is currently available against the virus, it is unarguably the largest economic threat to the global pig industry. ASFV is mainly replicates in the cytoplasm, but another form of nuclear replication is also detected in the early phase of the infection^19^. The ASFV genome is 190 kbp long and contains more than 190 open reading frames (ORFs), although the exact numbers of genes and proteins are unknown^20^. Approximately 20 viral genes are believed to participate in transcription and mRNA processing^21^, whereas at least 17 genes play a role in the replication, repair, and modification of DNA^22,23^. Depending on the strain, approximately 30-50 genes are involved in the evasion of immune surveillance and in encoding virulence and host range factors^22–24^.

The temporal regulation of ASFV gene expression appears to be similar to that in poxviruses^21,25^, in which four kinetic classes of genes have been described. The expression of immediate early and early genes precedes DNA replication, whereas the intermediate and late genes are generated subsequently to the onset of DNA replication^21^. To date, only a few ASFV genes have been transcriptionally characterized in details. ASFV mRNAs have 5’ cap structures and 3’ poly(A) tails added by the viral capping enzyme complex and the poly(A) polymerase, respectively^26^. ASFV is not easy to propagate, but it replicates relatively well in porcine primary alveolar macrophages (PAMs) *in vitro*, although the sensitivity of naïve PAM culture to ASFV infection varies batch by batch^27^.

To our knowledge, only a single attempt has been made to characterize the ASFV transcriptome using RNA-sequencing. However, this study did not generate sufficient data to construct a detailed transcriptional map of the virus^28^. To provide a much more detailed transcription map and to gain more information about the transcription dynamics of the virus, we performed multiplatform sequencing using both SRS and LRS techniques. The presented data set represent a key resource for studying the ASFV transcriptome at different time points after infection, and of the effect of infection on the host gene expression.

Regarding the SRS approach, the MiSeq instrument (Illumina) was used (Figure 1 shows the coverage depth), whereas we applied the MinION portable sequencer from Oxford Nanopore Technologies (ONT) for full-length sequencing. The random-primed SRS library was run on a single MiSeq v3 flow cell, whereas three different ONT libraries [direct RNA sequencing (dRNA-Seq), direct cDNA sequencing (dcDNA-Seq) and amplified cDNA sequencing) were sequenced on three individual flow cells. Altogether the three LRS experiments resulted in 17,122,822 sequencing reads (Table 1), of which 139,711 aligned to the viral genome (MN715134.1). The longest average read length was obtained using the dcDNA technique (1299 bp). The average length for the amplified approach ranged between 598-993 bp, whereas the dRNA-Seq resulted in an average read length of 953 bp (Table 1). More details about the length and quality of sequencing reads are presented in Table 1, Figure 2 and Supplementary Table 1. The quality data from Illumina sequencing is presented in Table 2.

**Figure 1.**
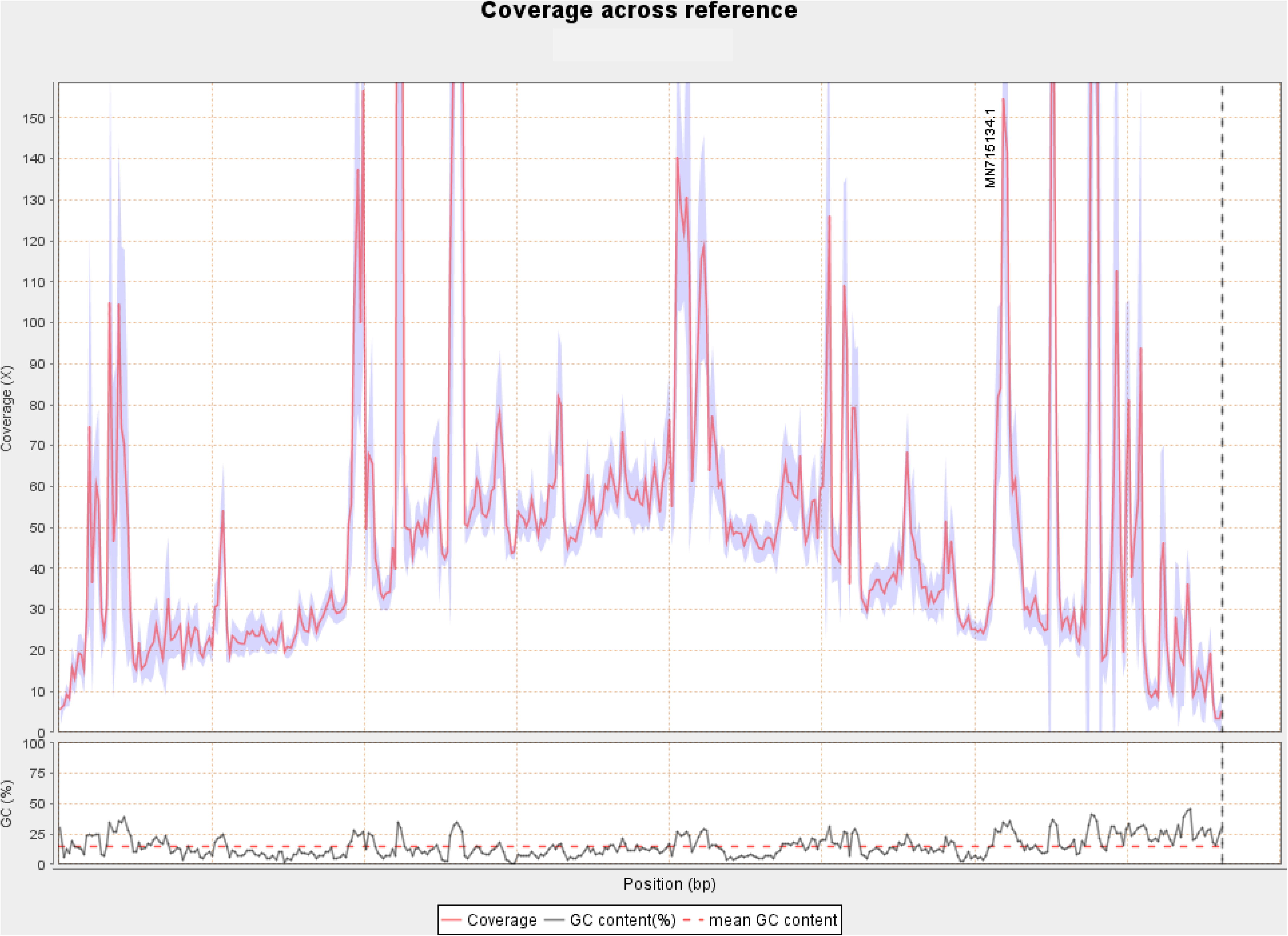
Illumina read depth along the African swine fever virus genome (MN715134.1). The G+C content of the virus is also shown at the bottom of the figure.

**Figure 2.**
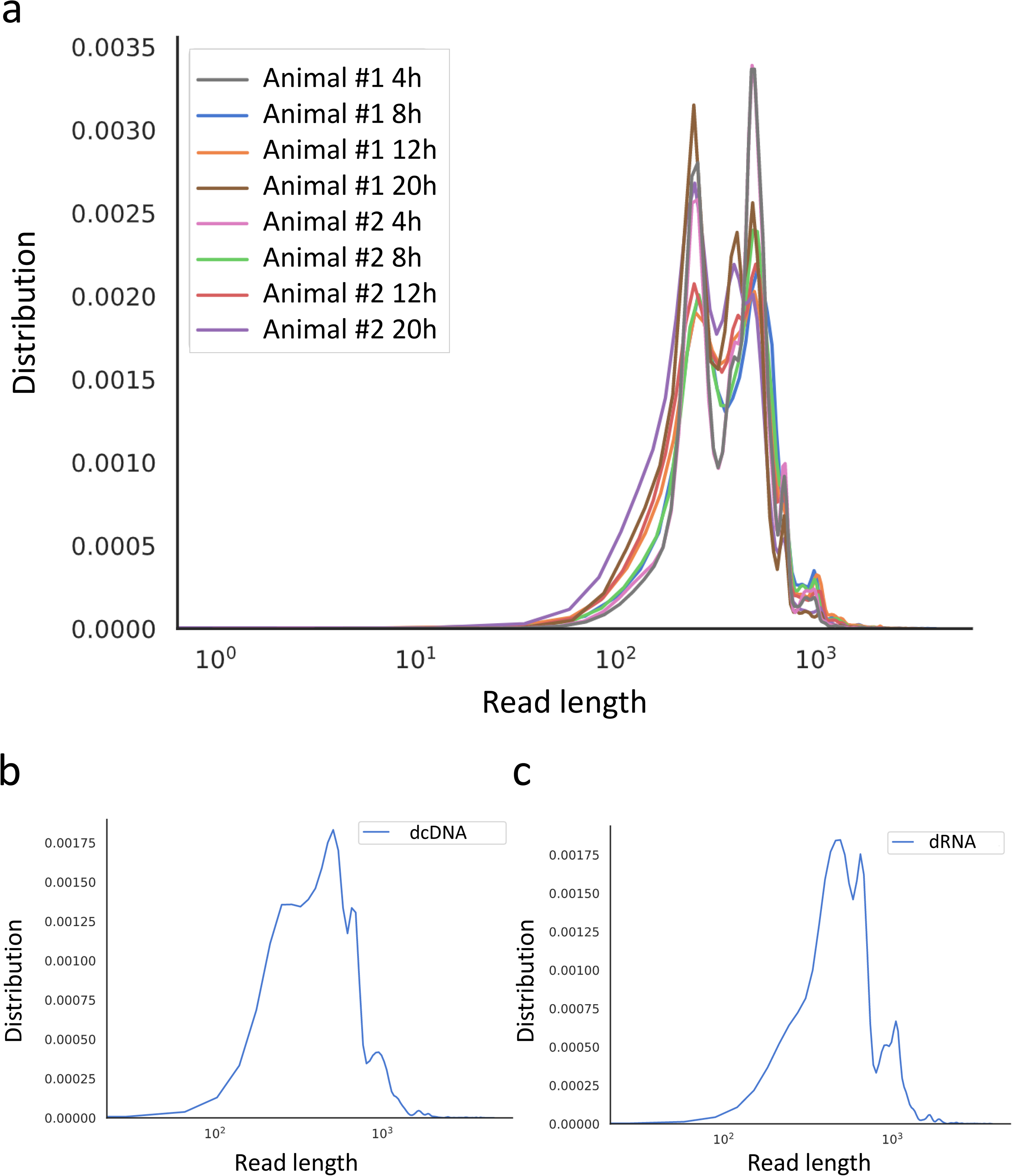
Aligned read length distribution. Line chart presentation of the average of aligned read lengths obtained via Nanopore sequencing. (a) Amplified cDNA sequencing at various time points. (b) Direct cDNA sequencing and (c) direct RNA sequencing using samples from multiple time points after the infection.

**Table 1.** Summary statistics of the Nanopore sequencing reads mapped to the viral genome for each run.

| Samples | Time point | Library preparation approach | Read count | Mapped read count | Mean read length |
| --- | --- | --- | --- | --- | --- |
| Animal #1 | 4 h | amplified cDNA | 1130020 | 18517 | 770,3±411,7 |
| Animal #1 | 8 h | amplified cDNA | 548671 | 9245 | 992,8±592,9 |
| Animal #1 | 12 h | amplified cDNA | 1230079 | 12779 | 899,4±514,1 |
| Animal #1 | 20 h | amplified cDNA | 2759917 | 33189 | 664,2±378,7 |
| Animal #2 | 4 h | amplified cDNA | 1562973 | 17668 | 764,9±426,4 |
| Animal #2 | 8 h | amplified cDNA | 978909 | 13346 | 961,8±548,4 |
| Animal #2 | 12 h | amplified cDNA | 1861829 | 13141 | 795,3±425,2 |
| Animal #2 | 20 h | amplified cDNA | 1567961 | 8878 | 597,9±365,5 |
| Animals #1 & #2 | mixed | dcDNA | 3591427 | 8587 | 1299,4±1045,4 |
| Animals #1 & #2 | mixed | dRNA | 1891036 | 4361 | 952,9±597,2 |

**Table 2.** Summary statistics of the Illumina reads aligning to ASFV genome.

|  |  |
| --- | --- |
| Mapped reads | 69 068 |
| Min read length | 58 bp |
| Max read length | 68 bp |
| Mean read length | 67,52 |
| Overlapping read pairs | 364 /1,05% |
| Duplicated reads (estimated) | 37 051/53,64% |
| Duplication rate | 36,12% |
| Clipped reads | 45 837/66,37% |
| Mean coverage | 50,1335 |
| St Dev mean coverage | 53,1682 |
| Mean mapping quality | 99,5 |
| Mean insert size | 445,27 |
| St Dev | 3 880,12 |
| P25/median/P75 | 223/274/346 |
| Insertions | 242 |
| Mapped reads with at least one insertion | 0,35% |
| Deletions | 396 |
| Mapped reads with at least one deletion | 0,57% |
| Homopolymer INDELS | 76,65% |

## Methods

The experimental design utilized in this study is shown in **Figure 3**. The applied reagents are listed in Table 3.

**Figure 3.**
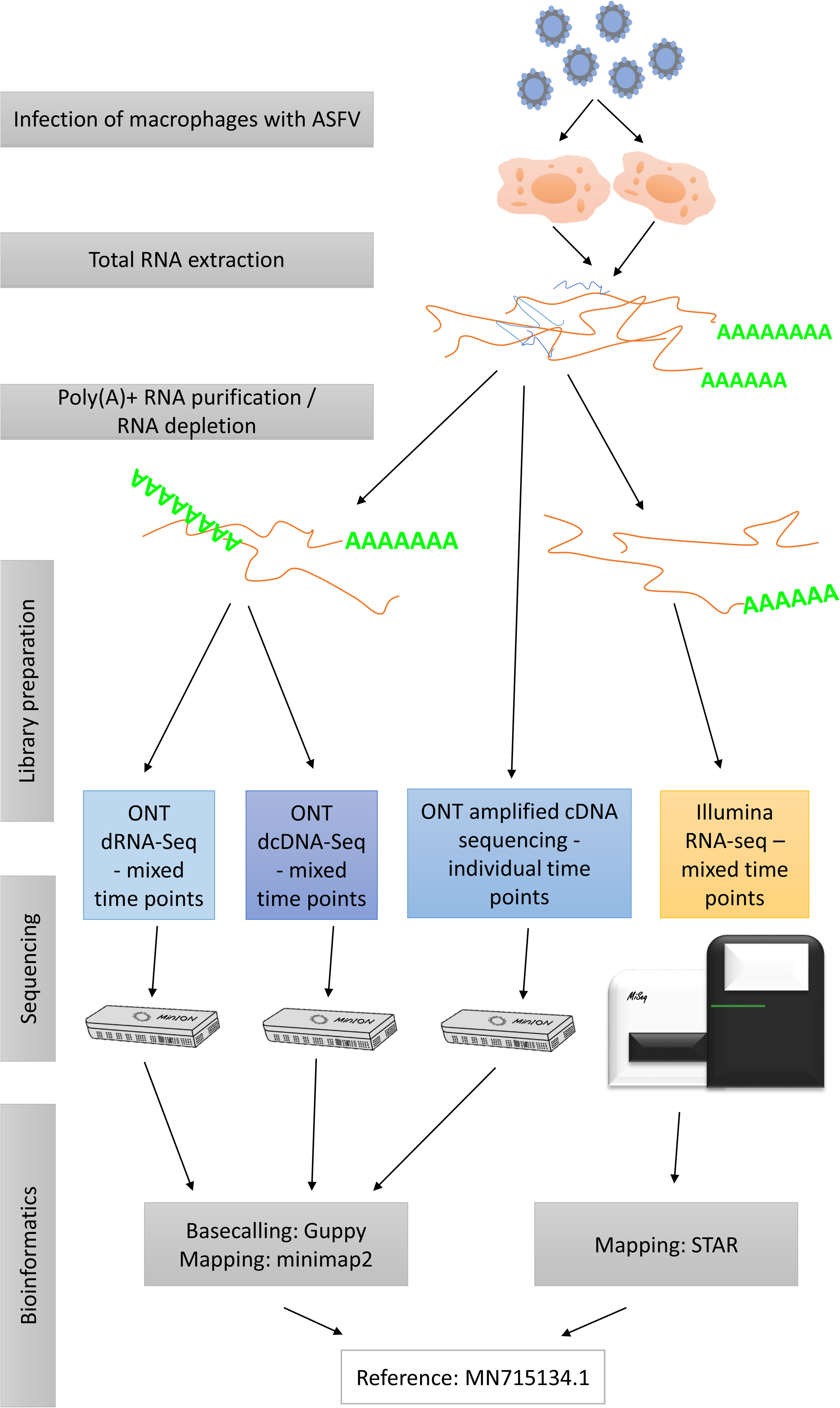
Flowchart diagram shows an overview of the methodological workflow of this study.

**Table 3.**
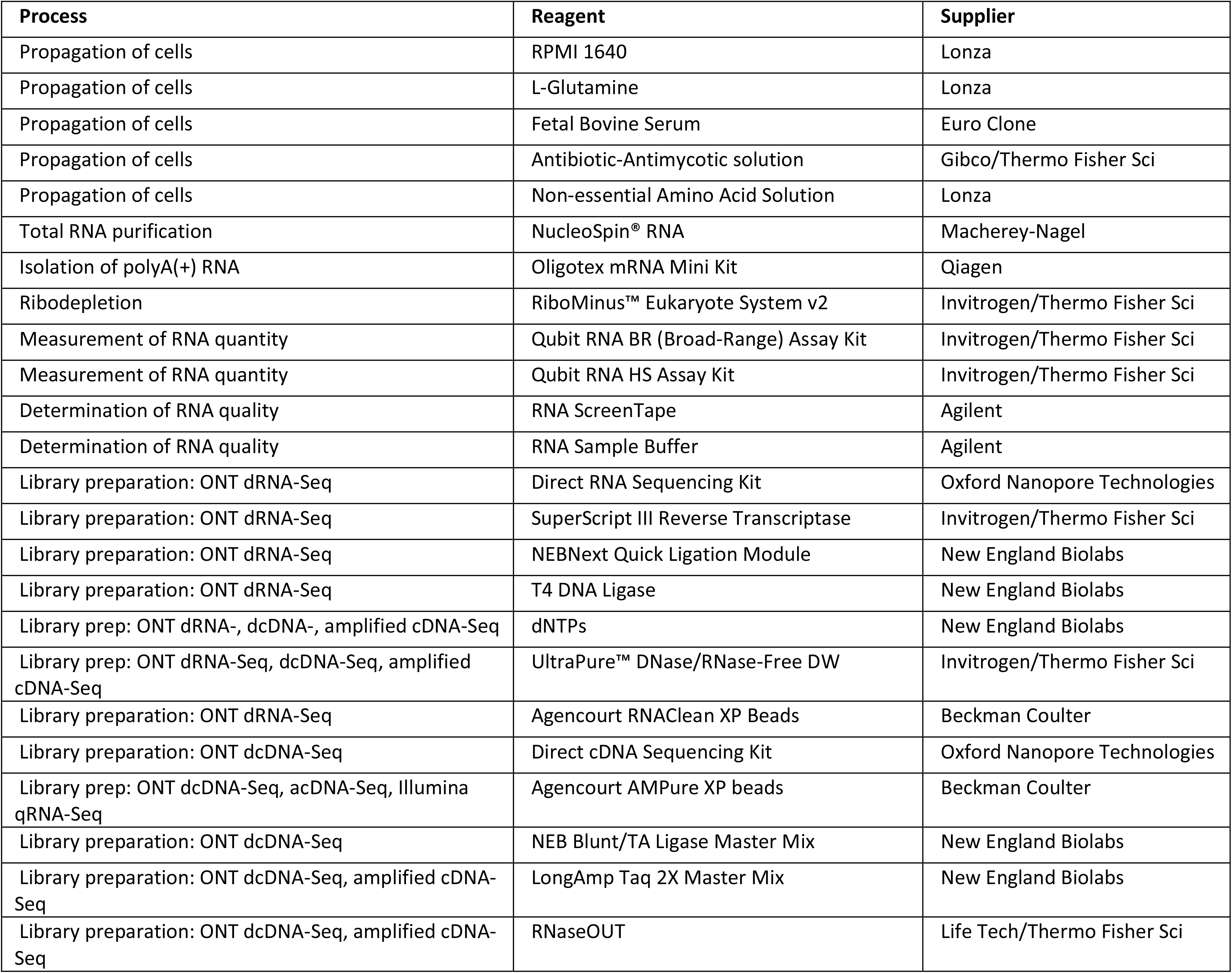
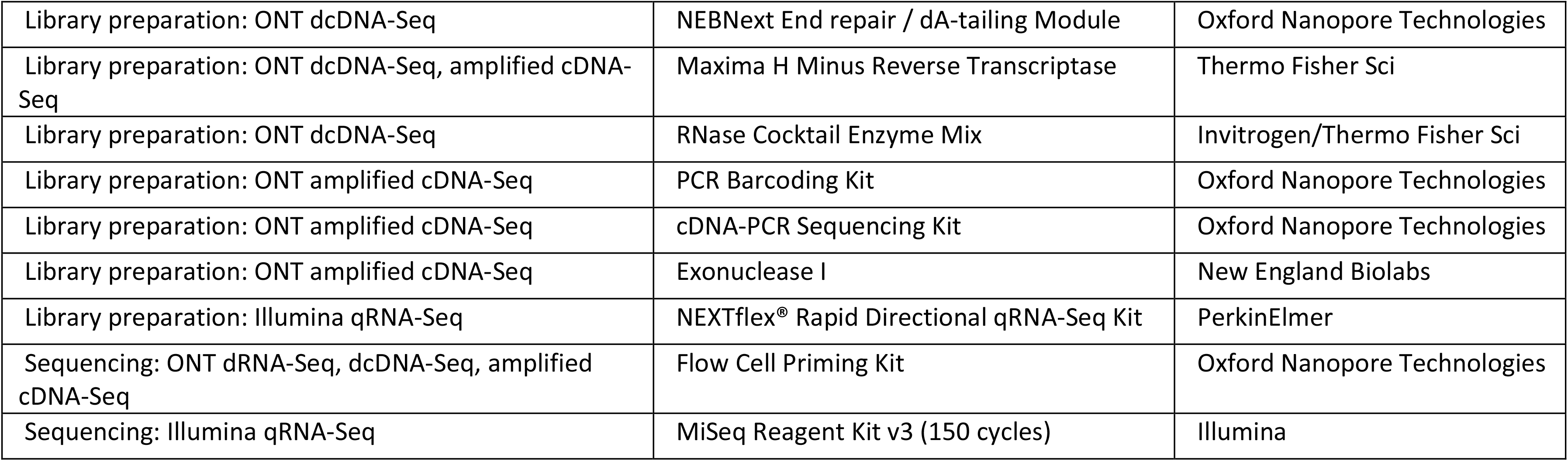
List of reagents and chemistries used in this work. Abbreviations: prep, preparation; DW, distilled water; Tech, technologies; ONT, Oxford Nanopore Technologies; Sci, Scientific; acDNA, amplified cDNA

### Cells and viruses

For the experiments fresh swine pulmonary macrophage (PAM) cells were harvested following the OIE Manual’s instructions (Manual). The cells were grown in RPMI 1640 containing L-glutamine (Lonza) medium supplemented with 10% fetal bovine serum (Euro Clone), 1% Na-pyruvate (Lonza), 1% antibiotic-antimycotic solution (Thermo Fisher Scientific), and 1% non-essential amino acid solution (Lonza) at 37°C in a humidified atmosphere containing 5% CO_2_. The highly virulent Hungarian ASFV isolate (ASFV_HU_2018 (ID Number: MN715134)) was used for infection. The infectious titer of the serially diluted viral stock was calculated in PAMs using an immunofluorescence (IF) assay as described previously^27^. Approval for the study of ASFV was obtained from the Ministry of Agriculture (Hungary), based on the 189 § (1) Chapter 3, of the 41/1997 (V. 28) Ministerial Decree, about the Veterinary Regulations (Approval number: ÉFHÁT/153-1/2019).

### Infection

PAMs were cultivated in 6-well plates at a density of 3.3 × 10^5^ cells/well and infected at a multiplicity of infection (MOI) of 10 at 4 h after cell seeding. Supernatant was replaced with fresh medium at 1 h post-infection (p.i.), and infected cells were harvested at 4, 8, 12, and 20 h p.i. Mock-infected control cells were also harvested at 20 h p.i.

The efficiency of infection was monitored using IF in an infected control well fixed at 20 h p.i. The infection rate of PAMs remained at approximately 20% despite the high viral titer (MOI = 10) used for infection. Based on our findings and previous experiments, achieving an infection rate exceeding 20% is difficult and incidental in naïve PAMs in the first infection cycle, even when using a high MOI.

### RNA purification

#### Isolation of total RNA

A NucleoSpin^®^ RNA (Macherey-Nagel) kit was used for RNA purification following the manufacturer’s instructions. In brief, the supernatant was removed from all wells of the 6-well plate, and 2 × 10^6^ cells were lysed with 400 µl of RA1 lysis buffer and 4 μl of β-mercapthoethanol solution. Then, the lysates were transferred to NucleoSpin filters. After centrifugation at 11,000 × *g* for 1 min, 350 μl of 70% ethanol were added to the lysates. The solutions were transferred to the columns, and after centrifugation at 11,000 × *g* for 30 s, the membranes were washed with 350 μl of MDB buffer. After repeated washing, 95 μl of a DNase reaction mixture (10 μl of reconstituted rDNase + 90 μl of reaction buffer) were added, and the membranes were incubated at room temperature for 15 min. The membranes were washed with 200 μl of RAW2 buffer and 650 μl of RA3 buffer, and the tubes were centrifuged at 11,000 × *g* for 30 s. Finally, 250 μl of RA3 buffer were added, and the tubes were centrifuged at 11,000 × *g* for 2 min. RNA was eluted with 60 μl of RNase-free H_2_O and centrifuged at 11,000 × *g* for 1 min. All buffers were supplied from the kit.

#### Purification of polyadenylated RNA

For the various Nanopore sequencing approaches, the polyA(+) RNA fraction of total RNA was isolated using the “Spin Columns method” from the Oligotex mRNA Mini Kit (Qiagen).

#### Ribosomal RNA depletion

For Illumina sequencing, ribosomal RNA (rRNA) was eliminated from the total RNA samples using the RiboMinus™ Eukaryote System v2 (Invitrogen/Thermo Fisher Scientific).

#### Determination of RNA quantity and integrity

RNA samples were quantified, and their quality was assessed using a Qubit 4 Fluorometer (Invitrogen/Thermo Fisher Scientific) with a Qubit RNA BR (Broad-Range) or Qubit RNA HS assay kit and with RNA Screen Tape and Sample buffer using TapeStation 4150 (Agilent). RNA with an RNA integrity number (RIN) exceeding 9.4 was used for cDNA generation (Figure 3). RNA samples were stored at −80°C until further use.

### Library preparation for Nanopore sequencing

#### Direct RNA sequencing – using samples from mixed time points

The dRNA-Seq method using a direct RNA sequencing kit (SQK-RNA002, Version: DRS_9080_v2_revK_14Aug2019) from ONT was used for amplification-free sequencing. This approach is highly recommended to explore special features of native RNA (e.g., modified bases) and avoid potential biases associated with reverse transcription (RT) or PCR. Total RNA from eight samples (two parallel experiments from 4, 8, 12, and 20 h p.i.) was mixed together, and then the polyA(+) fraction of RNA was purified from the sample mix. One hundred nanograms from the polyA-tailed RNA were diluted to 9 μl and then mixed with RT Adapter (oligo dT-containing T_10_ adapter), RNA CS (both from the Nanopore kit; the latter was used to monitor the sequencing quality), NEBNext Quick Ligation Reaction Buffer, and T4 DNA ligase (both from New England Biolabs). The mixture was incubated for 10 min at room temperature, and then RT was conducted to generate first-strand cDNA using the following protocol: 2 μl of 10 mM dNTPs (New England Biolabs), 5× first-strand buffer, 4 μl of 0.1 M DTT (both from Invitrogen SuperScript III) and 9 μl of UltraPure™ DNase/RNase-Free distilled water (Invitrogen/Thermo Fisher Scientific) were added, and then the sample was mixed with 2 μl of SuperScript III enzyme (Thermo Fisher Scientific) to obtain a 50-μl final reaction volume. RT was performed in a Veriti thermal cycler (Applied Biosystems) at 50°C for 50 min, and the reaction was subsequently terminated at 70°C for 10 min. RNA-cDNA hybrids were purified using Agencourt RNAClean XP Beads (Beckman Coulter; sample-bead ratio = 1:1.8), washed with freshly prepared 70% ethanol, and eluted in 20 μl of UltraPure™ nuclease-free water. The sample was then ligated to the RNA adapter (RMX from the ONT kit, 6 μl) at room temperature for 10 min using 8 μl of NEBNext Quick Ligation Reaction Buffer, 3 μl of T4 DNA ligase, and 3 μl of nuclease-free water. The ligation reaction was followed by a final purification step using 40 μl of XP Beads (sample:bead ratio = 1:1) to remove any potential remaining salts, proteins, or other contaminants. Samples were washed with wash buffer (ONT) and eluted in elution buffer (ONT). After the Qubit 4 Fluorometer measurement, 100 fmol from the library were loaded onto an R9.4 SpotON Flow Cell.

#### Direct cDNA-Seq – from mixed time points

Viral and host transcripts were also sequenced on a MinION sequencer following the instructions of the direct cDNA sequencing kit (SQK-DCS109; Version: DCS_9090_v109_revJ_14Aug2019; ONT). This protocol is based on strand switching, and it is highly optional for the generation of full-length cDNA for the identification of potential novel transcript isoforms without potential PCR bias. The starting material was 100 ng of a poly(A)+ RNA mixture from various time points of infection (4, 8, 12, and 20 h p.i.). An oligo dT-containing VN primer (VNP; 2.5 μl from the 2-μM stock) and 1 μl of dNTPs (10 μM) were added to the RNA. The total volume of the reaction was 11 μl. After 5 min of incubation at 65°C, the following components were added: 4 μl of 5× RT buffer (from the Maxima H Minus Reverse Transcriptase kit, Thermo Fisher Scientific), 1 μl of RNaseOUT™ (Thermo Fisher Scientific), 2 μl of strand switching primer (SSP, 10 μM) from the ONT kit and 1 μl of nuclease-free water. This mixture was pre-heated at 42°C for 2 min, and 1 μl of Maxima H Minus Reverse Transcriptase was added. RT was conducted at 42°C for 90 min, and finally, the reaction was stopped by incubation at 85°C for 5 min. One microliter of RNase Cocktail Enzyme Mix (Thermo Fisher Scientific) was used to degrade the RNA in the sample. Incubation was performed at 37°C for 10 min. Before the second-strand synthesis, the sample was cleaned using AMPure XP Beads (Beckman Coulter). Freshly prepared 70% alcohol was used to wash the samples, which were finally eluted in 23 μl of nuclease-free water. Then, 2× LongAmp Taq Master Mix (25 μl; New England Biolabs) was used to synthesize second-strand cDNA using 2 μl of the PR2 primer (ONT). Samples were incubated using the following “only one cycle protocol:” denaturation at 94°C for min, annealing at 50°C for 1 min, and elongation at 65°C for 15 min. The double-stranded cDNAs were purified using AMPure XP method (sample:bead ratio = 5:4) and eluted in 50 μl of nuclease-free water. NEBNext Ultra II End-prep reaction buffer (7 μl) and NEBNext Ultra II End-prep enzyme mix (3 μl, both from New England Biolabs) were added to each sample. This end repair process was performed at 20°C for 5 min, followed by a 5-min incubation at 65°C. Enzymes and buffers were removed from the reaction using the AMPure XP purification method, and then the 45-μl sample was subjected to adapter ligation. Five microliters from the ONT Adapter Mix (AMX) and 50 μl of Blunt/TA Ligation Master Mix (New England Biolabs) were mixed with each sample, and the mixture was incubated at room temperature for 10 min. A final AMPure XP purification was conducted to remove any excess proteins, nucleotides, and salts from the DNA library. Adapter Bead Binding Buffer (from the Nanopore kit) was used to wash the beads, and the library was eluted using elution buffer (part of the Nanopore kit). The samples were quantified using Qubit 4, and then 200 fmol from the sample were loaded on two MinION SpotON Flow Cells.

#### Amplified cDNA-sequencing – from different time points

Samples from each time point (4, 8, 12, and 20 h p.i.) were sequenced by the ONT MinION device using the cDNA-PCR Barcoding protocol (SQK-PCS109 and SQK-PBK004; Version: PCSB_9086_v109_revK_14Aug2019). This protocol is recommended to identify and quantify full-length transcripts, discover novel isoforms, and splice variants and fusion transcripts from a low amount of starting material (total RNA) to generate large amounts of cDNA data. Approximately 50 ng of each of the samples were used for library preparation. One microliter of VNP (2 μM) and 1 μl of dNTPs (10 μM) were added to the RNA (9 μl) and incubated at 65°C for 5 min. The strand-switching buffer mixture (4 μl of 5× RT buffer, 1 μl of RNaseOUT, 1 μl of nuclease-free water, and 2 μl of SSP) was added to the samples, which were incubated at 40°C for 2 min. RT was conducted by adding 1 μl of Maxima H Minus Reverse Transcriptase at 42°C for 90 min. The enzyme was inactivated by increasing the temperature to 85°C for 5 min. Twenty-five microliters of 2× LongAmp Taq Master Mix, 1.5 μl of one of the Low Input barcode primers (LWB01-12, from the ONT’s SQK-PBK004 kit, Table 4), and 18.5 μl of nuclease-free water were included in the RT reaction mixture. Table 5 shows the PCR conditions. PCR products were treated with 1 μl of exonuclease (New England Biolabs, 20 units), and the mixture was then incubated at 37°C for 15 min, followed by 80°C for 15 min. AMPure XP Beads was used for purification (sample:bead ratio = 5:4), and the clean sample was eluted in 12 μl of elution buffer.

**Table 4.**
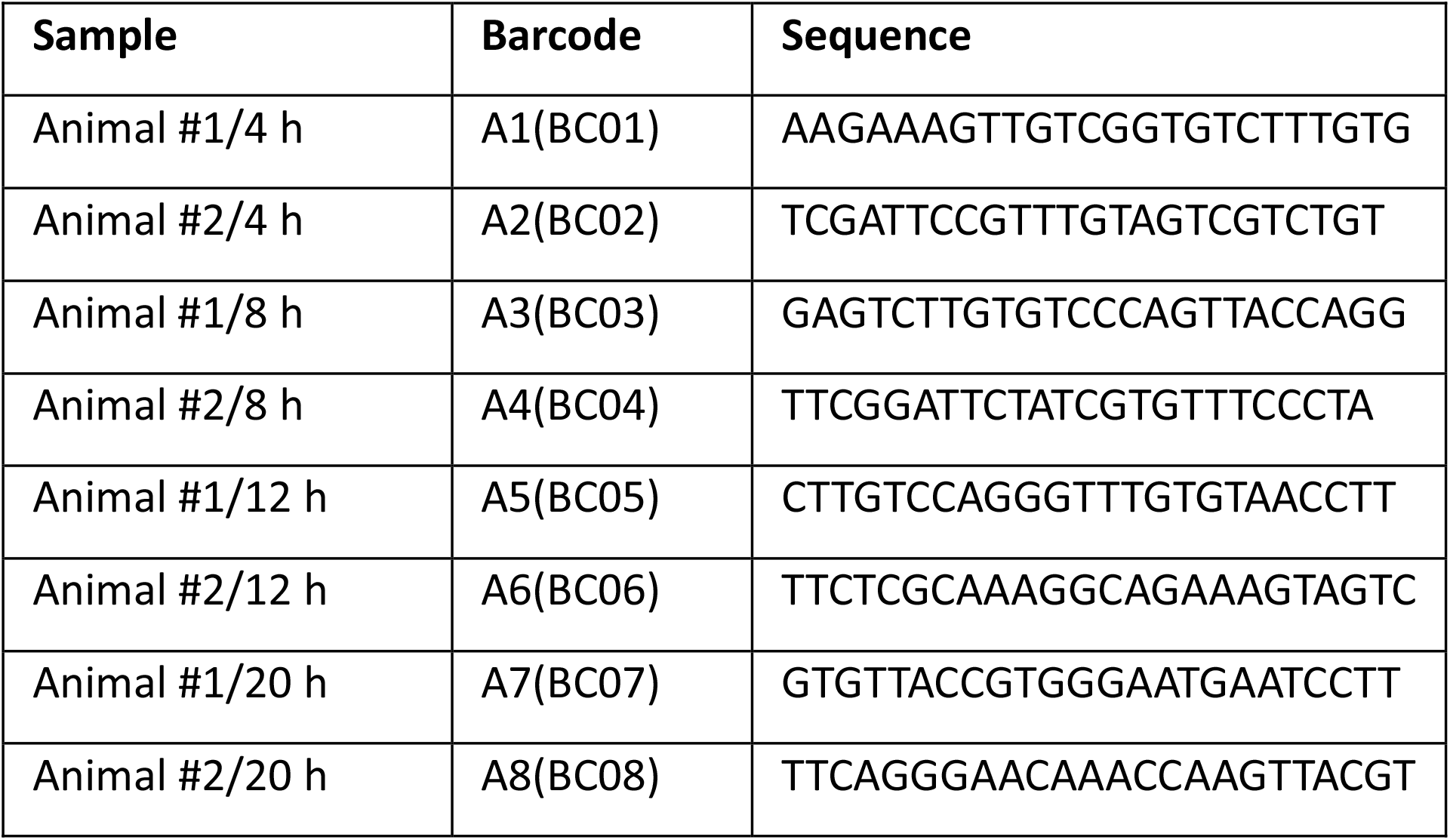
Barcode identifiers and sequences used for library preparation for the amplified cDNA sequencing approach.

**Table 5.** The protocol used for the amplification of the cDNA samples.

| Cycle step | Temperature | Time | No. of cycles |
| --- | --- | --- | --- |
| Initial denaturation | 95°C | 30 s | 1 |
| Denaturation | 95°C | 15 s | 14 |
| Annealing | 62°C | 15 s | 14 |
| Extension | 65°C | 4 min | 14 |
| Final extension | 65°C | 6 min | 1 |

### Library preparation for Illumina sequencing

A NEXTflex^®^ Rapid Directional qRNA-Seq Kit (PerkinElmer) was used to sequence the whole ASFV transcriptome via a conventional short-read approach. We used 25 ng (in 14 μl) of an rRNA-depleted RNA mixture (taken at 4, 8, 12, and 24 h p.i.) as the starting material. The first step was enzymatic fragmentation of the RNA using NEXTflex^®^ RNA Fragmentation Buffer (5 μl). The reaction was conducted at 95°C for 10 min followed by first-strand cDNA synthesis. First, 1 μl of NEXTflex^®^ First Strand Synthesis Primer was added to the reaction mixture, which was heated at 65°C for 5 min and then subsequently placed on ice. NEXTflex^®^ Directional First Strand Synthesis Buffer (4 μl) and NEXTflex^®^ Rapid Reverse Transcriptase (1 μl) were then added. RT was performed using the following program: incubation at 25°C for 10 min, heating at 50°C for 50 min, and termination at 72°C for 15 min. This step was followed directly by second-strand cDNA synthesis via the addition of 25 μl of NEXTflex^®^ Directional Second Strand Synthesis Mix (with dUTPs) at 16°C for 60 min. The product was cleaned using AMPure XP Beads (sample:bead ratio = 5:9). The bead-sample mix was washed with 80% alcohol, and resuspension buffer (from the NEXTflex^®^ Kit) was used for the final elution. Polyadenylation of the double-stranded cDNAs was performed using 4.5 μl of NEXTflex^®^ Adenylation Mix at 37°C for 30 min. The reaction was terminated by heating at 70°C for 5 min. Molecular Index Adapters (2 μl, 1 μM from the NEXTflex^®^ Kit) were ligated to the sample at 30°C (10 min) using the NEXTflex^®^ Ligation Mix (27.5 μl). Prior to amplification, each sample was washed with AMPure Beads. First, 1 μl of NEXTflex^®^ Uracil DNA Glycosylase was mixed with the sample, which was incubated at 37°C for 30 min, followed by heating at 98°C for 2 min. The sample was placed on ice, and the following components were added: 12 μl of PCR Master Mix, 2 μl of qRNA-Seq Universal forward primer, and 2 μl of qRNA-Seq Barcoded Primer (sequence: AACGCCAT; all were provided with the kit). The samples were amplified according to the protocol summarized in Table 6. The PCR products were washed with AMPure XP Beads, which was followed by a second purification.

**Table 6.** Table shows the PCR parameters used for Illumina library preparation.

| Cycle step | Temperature | Time | No. of cycles |
| --- | --- | --- | --- |
| Initial denaturation | 98°C | 2 min | 1 |
| Denaturation | 98°C | 30 s | 15 |
| Annealing | 65°C | 30 s | 15 |
| Extension | 72°C | 60 s | 15 |
| Final extension | 72°C | 4 min | 1 |

### Quantification and validation of the Illumina library

A Qubit 4 Fluorometer was used for the concentration measurement, whereas an Agilent TapeStation 4150 was used for the quality analysis (Figure 4).

**Figure 4.**
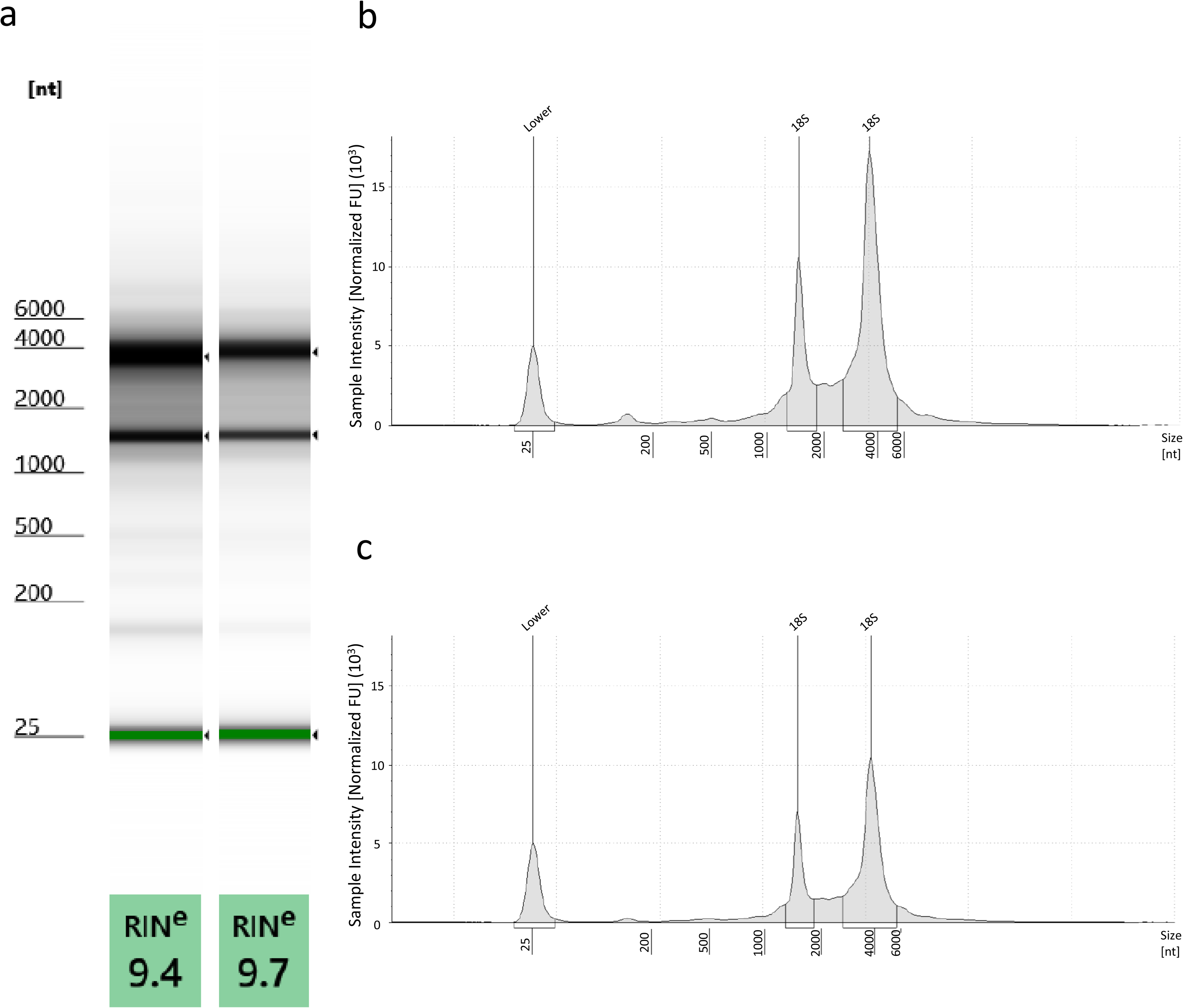
Total RNA analysis was conducted using the TapeStation 4150 system. High quality [RNA integrity number (RIN) = 9.4-9.7]. RNA samples were used for sequencing. (a) ScreenTape gel image shows the separation profile of two total RNA samples (Animal #1 8 h and Animal #2 12 h) along with the RINs. Panels (b) and (c) show representative electropherograms of total RNA from Animal #1 8 h and Animal #2 12 h, respectively. The electropherograms show ribosomal 28S and 18S RNA peaks and the lower marker.

### Sequencing on the Illumina MiSeq instrument

The sequencing-ready library (12 pM) was loaded onto a flow cell from Illumina MiSeq Reagent Kit v3 (150-cycle format) and sequenced on a MiSeq sequencer.

### Read processing

#### ONT sequencing

Guppy software v3.4.5 (ONT) was used for base calling of the data from MinION sequencing. The raw reads were aligned to the ASFV reference genome (NCBI Nucleotide accession: MN715134.1) using the minimap2 software suite^29^ with the following options: -ax splice -Y -C5 --cs. In-house scripts were used to obtain the quality information presented in this study.

#### Illumina sequencing

Raw reads were trimmed using Cutadapt software^30^, the aforementioned viral reference genome was indexed using STAR aligner v2.7.3a^31^ with the following settings: --genomeSAindexNbases 8, followed by the mapping of the reads with default options. Samtools^32^ was used to sort the sam files and to generate and index the bam files. The Qualimap v2.2.1 application^33^ was used to generate quality information from the Illumina dataset.

## Data Records

All data (Data Citation 1) have been uploaded to the European Nucleotide Archive under the accession number PRJEB36723. All reads were mapped to the MN715134.1 genome. All data can be used without restrictions.

## Technical Validation

The amounts of purified total RNA, polyA(+) RNA, and rRNA-depleted samples; generated cDNAs; and final sequencing libraries were quantified by a Qubit 4 Fluorometer using Qubit RNA Broad-Range, High Sensitivity RNA, and High Sensitivity dsDNA Assay Kits. The Agilent TapeStation 4150 system was applied to detect the integrity of total RNA and perform a quality check of the Illumina libraries. In the present study, RNAs samples with RIN ≥ 9.4 were subjected to construct the sequencing libraries (Figure 4).

## Usage Notes

To the best of our knowledge, there are no data on the ASFV transcriptome; therefore, this dataset was primarily generated to characterize the RNA profile of the virus. The dataset can be used to detect RNA isoforms, including length (alternative 3’ and 5’) variants and monocistronic, bicistronic, polycistronic, and complex transcripts, and to discover transcriptional overlaps and the complexity of the genetic regulation of ASFV. The Nanopore dataset allows a time-course evaluation of the full-length transcriptomes of both the virus and host. The published binary alignment (BAM) files contain the reads mapped to the MN715134.1 ASFV reference genome. The BAMs (using samtools and bedtools^34^) can be converted to FastQ files, which extend the potential usage of the data; e.g., they can be aligned to the host genome. The BAM files can be analyzed using various bioinformatics tools, such as samtools, bedtools, or the Genome Analysis Toolkit^35^. The Nanopore data generated with different library preparation approaches can be compared to analyze the differences between the sequencing chemistries, as well as the effect of RT and PCR reactions on the length and quality of the reads. The provided dataset is also useful for comparing the performance of the utilized sequencing platforms. The Tombo tool^36^ can be used to identify CpG methylation patterns and base modifications (e.g., A to I editing) from raw (fast5) Nanopore sequencing data, or the EpiNano^37^ algorithm can be applied to detect potential m6A RNA modifications. The dataset can be further analyzed using various bioinformatics program packages (e.g., bedtools, samtools) or visualized using software packages such as the Integrative Genomics Viewer^38^, Savant Genome Browser^39^, and Geneious^40^.

## Code Availability

1. Guppy v3.4.5: https://community.nanoporetech.com/downloads?fbclid=IwAR2IchRL4gDnfA6h996UkN4vS5pbBu6rUtKVFX3aTiBHsWFknglQ6FyvPkg (Available for Nanopore Community members, free registration here: https://nanoporetech.com/community#register&modal=register)
2. minimap2: https://github.com/lh3/minimap2
3. STAR: https://github.com/alexdobin/STAR
4. samtools: https://github.com/samtools/samtools

## Acknowledgments

This study was supported by the National Research, Development, and Innovation Office grants FK 128252 to DT and K 128247 to ZB. The publications cost was covered by the University of Szeged Open Access Fund (Grant no: 4604).

## Author contributions

F.O. participated in the infection experiments, and the purification of the total RNA from the virus, and with writing the manuscript.

D.T. performed the MinION sequencing (dynamic dataset) and the Illumina sequencing, participated in data analysis and wrote the manuscript.

G.T. generated the ONT dRNA sequencing libraries and conducted data handling and processing.

Z.C. conducted MinION direct cDNA sequencing and the Illumina sequencing.

N.M. carried out bioinformatics analysis.

A.D. performed the isolation of poly(A)+ RNAs and the rRNA removal from total RNAs.

I.P. participated in the purification of poly(A)+ RNA and data handling.

I.M. conducted the infection experiments, purified the viral total RNA samples

T.M. prepared the pulmonary macrophages

T.V. participated in the infection experiments, and the purification of the total RNA from the virus

Z.Z. designed the research plan and wrote the manuscript

Z.B. conceived and designed the experiments, managed the study, and wrote the manuscript. All authors read and approved the final paper.

## Supplementary material

**Supplementary Table 1. Detailed data of the sequencing reads that mapped to the African swine fever virus genome**.

